# Synonymous codon usage is biased for and against m^6^A RRACH motifs in mammals

**DOI:** 10.64898/2026.07.09.737571

**Authors:** Leo D. Creasey, Eran Tauber

## Abstract

RNA methylation at N^6^-adenosine (m^6^A) predominantly occurs within RRACH motifs, yet the forces shaping these motifs in coding regions remain unclear. Here we show that the synonymous codon combinations able to form or disrupt RRACH sites are used non-randomly across mammals. Using 13,491 protein-coding genes from 261 species, we identified genes significantly enriched or depleted in RRACH motifs, a pattern consistent with gene-specific selection for or against m^6^A potential. Genes enriched in RRACH sites were linked to ubiquitin-like conjugation and cell cycle regulation, whereas transmembrane and HOX genes were RRACH-poor, likely reflecting sequence incompatibility with CpG dinucleotides. Cross-species comparison with *Caenorhabditis elegans*, which lacks mRNA m^6^A methylation, revealed reciprocal RRACH frequencies, as expected if these motifs are under selection in m^6^A-competent genomes but evolve without this constraint otherwise. At the codon level, specific amino acid pairs, particularly threonine-ending dyads, were biased toward RRACH-forming codons while others were depleted, indicating that synonymous codon choice is skewed for and against motif formation. RRACH motifs were also non-randomly distributed along coding sequences, depleted near start codons and enriched toward the 3′ end, consistent with known m^6^A profiles. Finally, analysis of cancer mutations revealed tissue-specific gain and loss of RRACH sites, reflecting context-dependent remodeling of methylation potential. Together, these results show that synonymous codon usage is systematically biased for and against m^6^A RRACH motifs, pointing to an evolutionary coupling between the genetic code and the epitranscriptomic landscape.

## 1 Introduction

RNA modification has been detected in over 170 different forms (Tang et al., 2023), but the most prevalent is RNA methylation in the form of methyl-6-Adenine (m6A). m6A is found throughout large swathes of the animal kingdom, from yeast and bacteria (Bodi et al., 2010; Deng et al., 2015), to mammals (Song et al., 2021), and has been implicated in many different biological processes, such as mRNA translation and stabilization (Wang et al., 2015), mRNA decay (Wang et al., 2014), and alternative splicing (Pendleton et al., 2017). m6A occurs in an estimated 20-60% of transcripts, and is more frequently found near the 3’ end of mRNA transcripts (Meyer et al., 2012). Further, RNA m6A methylation is not restricted to mRNA, and has been found in tRNA, rRNA, and long non-coding RNA (Hori, 2014; Sergiev et al., 2018). m6A methylation most commonly occurs on RRA_m_CH sites (R = A or G, H = A, C, or U). An important feature of m6A methylation is that it is reversible (Fu et al., 2014), which has given rise to a set of proteins that add methylation (writers), remove methylation (erasers), and detect methylation leading to biological action (readers). Of these, the most prominent writers are methyltransferase-like 3 (METTL3), METTL14, and Wilms tumor 1-associated protein (WTAP). The proteins Fat mass and obesity-associated (FTO) and alkB homolog 5 (ALKBH5) are the core human methylation erasers, and YT521-B homology domain containing proteins (YTHDF1, YTHDF2, YTHDF3) are the most prominent readers that recognize m6A methylation (reviewed by Zaccara et al. 2019).

Dysregulation of RNA methylation has also been noted in many disease pathologies, with the most notable being cancer, where the versatility of m6A methylation means that genes linked with RNA methylation can either be upregulated or downregulated depending on the cancer type or organ (reviewed by Xue et al., 2022). For example, Hou et al.,(2020) demonstrated that upregulated METTL3 was found in a variety of digestive tract cancers, but in renal carcinomas, METTL3 was found to be downregulated (Li et al., 2017). A similar phenomenon can be observed in erasers such as FTO, which have been found to be upregulated in esophageal cancer (Liu et al., 2020) and downregulated in bladder cancer (Wen et al., 2020).

We have previously demonstrated a pro-epigenetic selection process that favors combinations of consecutive codons in coding sequences that result in formation of DNA methylation sites (CpG), across a large number of mammalian species (Creasey & Tauber, 2024). We coined these pairs of codons ‘CpG Codon Dyads’. We have also identified coding sequences in which the occurrences of CpG dyads were depleted, reducing CpG sites available for DNA methylation. Here, we sought to test whether codon usage may also be driven by selection for RNA methylation by the formation of RRACH sites in coding sequences. The structural flexibility and sequence degeneracy of RRACH motifs provide more opportunities for mutations than CpG sites, allowing these sequences to be more readily created or lost. Potentially, there are 48 codon pairs that may form an RRACH motif (Figure 1). In addition to these RRACH ‘dyads’, the RRACH motif may also be formed by codon ‘triads’ (NNR-RAC-HNN). Here, we investigated whether these sites are conserved, whether specific gene groups show an over- or under-abundance of RRACH sites, and whether particular amino acid pairings or triplets are overrepresented in RRACH motifs. Furthermore, we examined whether somatic mutations identified in cancer genomes generate more or fewer RRACH sites on average across cancer types.

**Figure 1.**
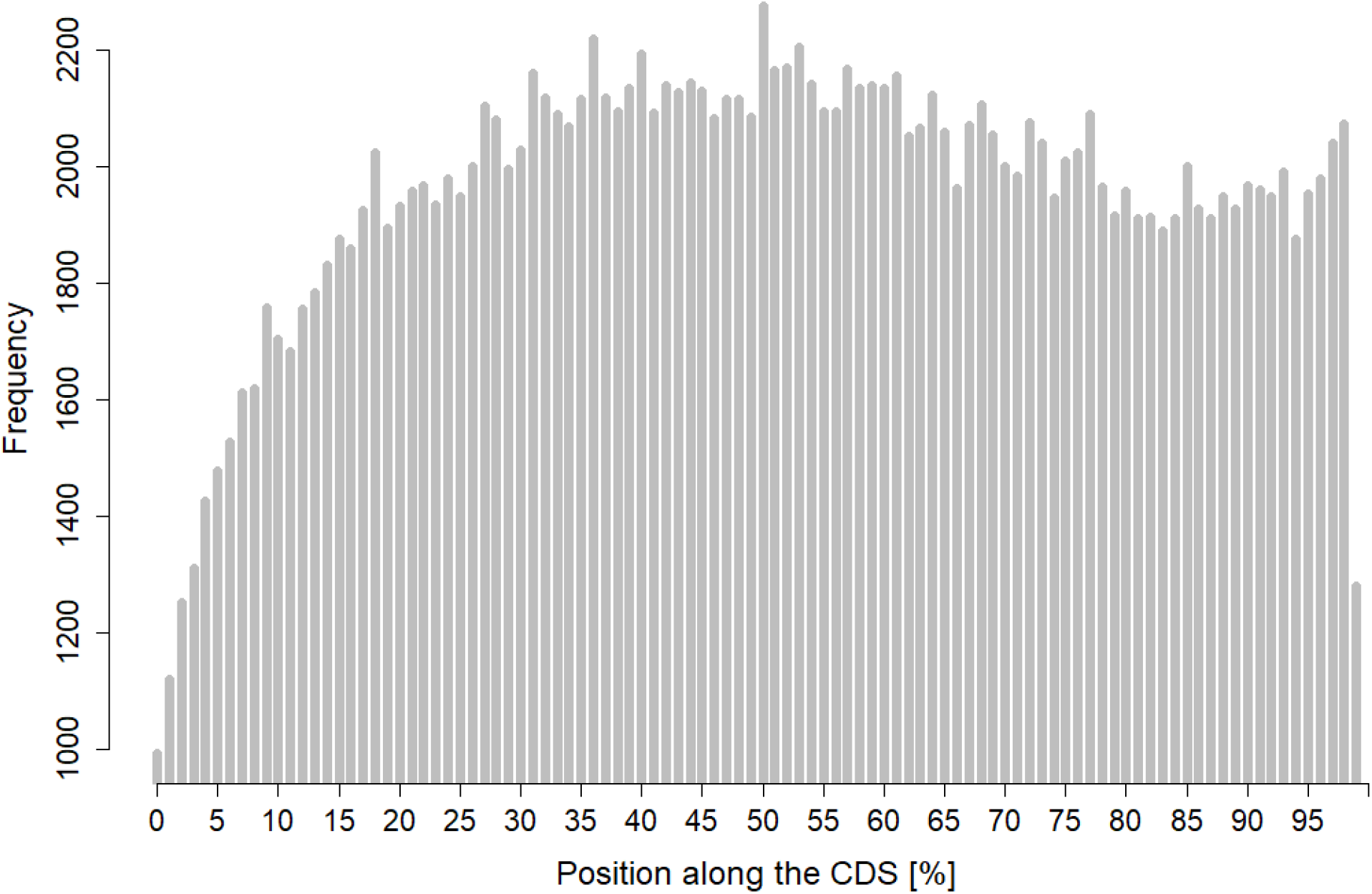
Frequency of RRACH sites along the relative length of coding sequences (CDS). The analysis is based on 13,491 protein-coding genes from 261 mammalian species.

## 2 Methods and Materials

Sequence data were obtained from Bowman et al. (2023) and consisted of alignments of 261 mammalian sequences across 13,491 protein-coding genes. A custom Python script was used to calculate, for each species and each gene, the number and positions of RRACH sites, enabling estimation of RRACH site conservation across species. The number of RRACH sites was normalized both for the number of species represented per gene (as some genes were absent in certain species) and for gene length (RRACH sites per number of base pairs). An average number of RRACH sites per sequence was also calculated. An RRACH site was considered conserved if ≥200 of the 261 species (76%) contained a RRACH codon dyad at that position.

Gene Ontology (GO) enrichment analysis was performed using the DAVID Bioinformatics Resources (Huang et al., 2009) to identify gene groups that were over- or under-represented in terms of RRACH site counts.

A separate Python script was developed to identify amino acid dyads and triads that could contain RRACH motifs. The script tracked the number of such motifs that did and did not contain RRACH sites, yielding observed frequencies, which were compared to expected frequencies using a χ^2^ (chi-squared) test performed in R (R Core Team, 2024).

In house code was developed to record GC and GC3 content of the human version of the coding sequences. Combined with the length of the gene, GC and GC3 ratios were generated on a per gene basis. Using in a Poisson GLM regression analysis, we determined if GC or GC3 ratio influenced the RRACH count across our genes.

Data on experimentally validated RNA methylation sites were obtained from RMBase v3.0 (Xuan et al., 2024). Custom code parsed these datasets to count methylated m^6^A sites within each gene’s coding sequence, from which the proportion of RRACH sites that were methylated was calculated and visualized in R.

A list of Caenorhabditis elegans orthologous genes was obtained using the online BioMart tools (Kinsella et al., 2011). Full C. elegans coding sequences matching the list of orthologues were retrieved from EMBL (Dyer et al., 2025). In-house code was used to identify orthologs within the dataset with high orthology confidence, as defined by BioMart, and to retrieve the corresponding FASTA sequences. The code then calculated both the frequency and density (frequency per gene length) of RRACH sites for each orthologous gene pair. In total, 1,783 orthologous gene pairs were analyzed. The data were subsequently subset in R to group the 500 genes with the highest and the 500 genes with the lowest RRACH density in the human sequences. Differences in RRACH density between the human and C. elegans orthologs were tested using the Wilcoxon rank-sum test.

To calculate amino acid dyad frequencies, custom Python code analyzed all human coding sequences to determine the frequency of each nucleotide dyad and its corresponding amino acid pair. For each amino acid dyad capable of forming an RRACH motif, the number of observed RRACH-containing dyads and the number of possible dyad combinations that could generate a RRACH motif were calculated. The normalized frequency was computed as the number of RRACH sites divided by the number of unique dyad sequences capable of generating RRACH.

This analysis was repeated for dyads that did not contain an RRACH site. All graphs were generated in R, and χ^2^ tests were implemented in Python. All custom scripts used in this study are available at https://github.com/Leo-Creasey/RNA_methylation_project.git.

Cancer mutation data were obtained from the Catalogue of Somatic Mutations in Cancer (COSMIC) (Sondka et al., 2024). Custom Python code compiled lists of mutations associated with each gene, stratified by the organ of origin. Two COSMIC datasets were analyzed: (1) mutations in known cancer-related genes and (2) a larger dataset encompassing all mutations observed in cancer samples.

Statistical analyses and data visualization were conducted in R, using the packages *ggplot2, viridis, gridExtra, tidyr*, and *dplyr* (Auguie, 2017; Garnier et al., 2024; Wickham, 2016; Wickham et al., 2023, 2024).

## 3 Results

### 3.1 Functional groups associating with either high or low abundance of RRACH sites

RRACH sites were analyzed across 13,491 protein-coding genes from 261 eutherian mammal species. We performed an overrepresentation analysis on the top 1,000 genes with the highest density of RRACH sites (approximately the top 7.4% most enriched genes). The results revealed several over-represented functional categories among genes with an abundance of RRACH motifs (Table 1). These were primarily associated with ubiquitin-like conjugation (enrichment score = 30.03) and cell cycle/division (enrichment score = 9.58). Additional, though less enriched, categories included centromere and kinetochore functions.

**Table 1.** GO functional annotation of genes enriched by RRACH sites.

| Cluster | ES | Category | Associated Terms | p-value | Genes (#) |
| --- | --- | --- | --- | --- | --- |
| 1 | 30.03 | UP_SEQ_FEATURE | Glycyl lysine Isopeptide | 2.7E-34 | 165 |
|  |  | UP_KW_PTM | Isopeptide bond | 6.1E-33 | 215 |
|  |  |  | Ubl conjugation | 5.0E-25 | 254 |
| 2 | 9.58 | UP_KW_BIOLOGICAL_PROCESS | Cell cycle | 1.7E-14 | 83 |
|  |  | GOTERM_BP_DIRECT | Cell division | 3.5E-10 | 48 |
|  |  | UP_KW_BIOLOGICAL_PROCESS | Mitosis | 2.5E-7 | 37 |
| 3 | 5.66 | GOTERM_MF_DIRECT | Microtubule binding | 1.6E-8 | 38 |
|  |  | UP_KW_CELLULAR_COMPONENT | Microtubule | 9.3E-7 | 36 |
| 4 | 5.17 | UP_KW_CELLULAR_COMPONENT | Centromere | 1.8E-7 | 25 |
|  |  | GOTERM_CC_DIRECT | Kinetochore | 1.1E-5 | 22 |
| 5 | 3.84 | UP_KW_BIOLOGICAL_PROCESS | DNA damage | 2.4E-5 | 43 |
|  |  |  | DNA repair | 9.3E-5 | 36 |

ES = Enrichment score produced by Functional Annotation Clustering in DAVID. Category Terms Defined: UP\_KW = Uniprot Keywords; GOTERM MF DIR = GO Term for Direct Involvement in Molecular Function; GOTERM BP\_DIR = GO Term for Direct Involvement in Biological Process; SMART protein domain database

**Table 2.** GO functional annotation of genes depleted by RRACH sites.

| Cluster | ES | Category | Associated Terms | p-value | Genes (#) |
| --- | --- | --- | --- | --- | --- |
| 1 | 12.93 | UP_KW_DOMAIN | Transmembrane helix | 1.6E-22 | 418 |
|  |  |  | Transmembrane | 1.6E-22 | 421 |
|  |  | UP_KW_CELLULAR_COMPONENT | Membrane | 8.0E-12 | 507 |
|  |  |  | Cell membrane | 1.4E-3 | 238 |
| 2 | 5.62 | INTERPRO | Homeobox CS | 2.5E-9 | 32 |
|  |  | SMART | HOX | 5.4E-8 | 32 |
|  |  | UP_SEQ_FEATURE | Homeobox | 4.5E-7 | 31 |
| 3 | 5.25 | KEGG_PATHWAY | Neuroactive ligand-receptor interaction | 5.7E-12 | 49 |
|  |  | UP_SEQ_FEATURE | Transmembrane Helical | 2.8E-8 | 84 |
|  |  | UP_KW_MOLECULAR_FUNCTION | Transducer | 6.0E-6 | 71 |
| 4 | 4.27 | INTERPRO<br>GOTERM_BP_DIRECT | EMP<br>Bleb assembly | 4.1E-6<br>4.5E-6 | 7<br>6 |
| 5 | 3.82 | SMART<br>GOTERM_MF_DIRECT | GCPR<br>Neuropeptide binding | 1.3E-5<br>4.7E-4 | 18<br>8 |

When the analysis was extended to include the 1,000 genes with the lowest RRACH density, we observed significant over-representation of transmembrane proteins (enrichment score = 12.93). Notably, HOX and other homeobox-containing genes were also enriched in this group (enrichment score = 5.62). Interestingly, our previous work demonstrated that these genes tend to contain a high abundance of CpG codon dyads (Creasey & Tauber, 2024). Given the mutual exclusivity of CpG and RRACH sequence motifs, this may explain the observed pattern. The analysis was repeated with top and bottom 500 genes, with similar results found (data not shown).

### 3.2 Distribution of RRACH sites along the coding sequence

Previous studies have shown that methylated RRACH sites are more frequently located near the 3′ end of coding sequences (Meyer et al., 2012). To determine whether RRACH sites in our dataset follow a similar pattern, we quantified the frequency of conserved RRACH sites relative to their position along the coding sequence (CDS) (Figure 1).

We found a relative depletion of RRACH sites near the start of the CDS, which leveled off approximately 30% along the length of the gene. A smaller peak was observed near the 3′ end, consistent with previous reports of RNA methylation enrichment in this region.

### 3.3 Effect of GC and GC3 content on RRACH frequency

GC content is a highly influential and selectively constrained feature of coding sequences, suggesting that it could potentially affect the frequency of RRACH sites. To test this hypothesis, we quantified GC content and GC3 content, defined as the GC proportion at the third synonymous codon position, and evaluated their association with RRACH frequency using Poisson generalized linear models.

GC content was significantly associated with RRACH frequency (p < 2 × 10^−16^); however, the effect size was minimal (-0.016). This corresponds to an expected 1.6% decrease in RRACH site frequency across the full range of GC content from 0% to 100%. A similar result was obtained for GC3, which also exhibited a statistically significant but very small effect (effect size = -0.0079, p < 2 × 10^−16^).

These findings suggest that, although GC content is significantly associated with RRACH frequency, its effect size is too small to be considered biologically meaningful.

### 3.4 Actual methylation rates of RRACH sites

Using the RMBase dataset, we estimated the number of methylated RRACH sites per gene and expressed this as a percentage of predicted RRACH sites *(number of m6A sites/number of predicted RRACH sites)×100*. The distribution of percentage methylation for each gene is shown in Figure 2.

**Figure 2.**
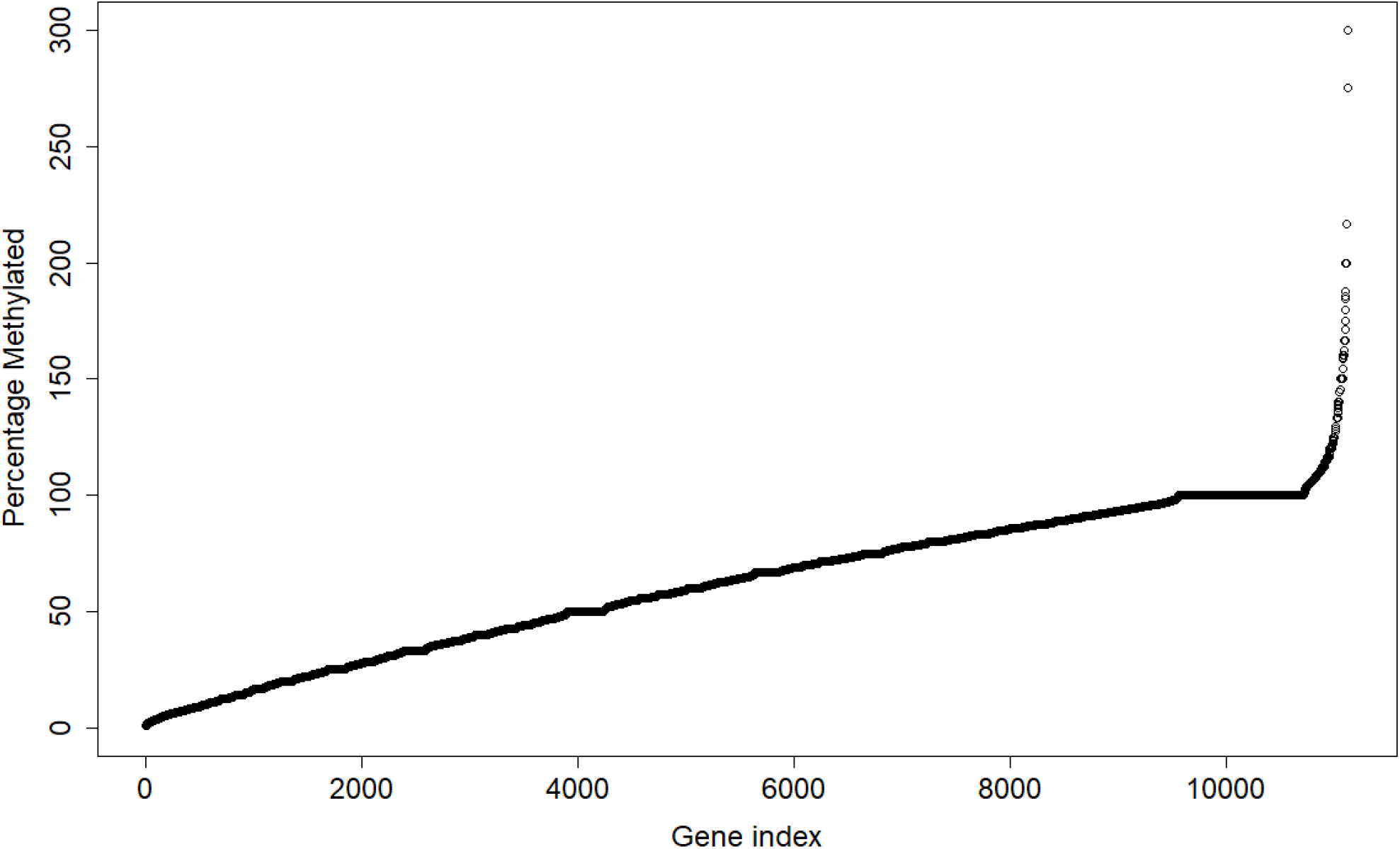
Percentage of RRACH sites that are methylated (n = 11,135). A limit was applied to the y-axis to improve data visualization; 14 data points were excluded as a result.

Across all genes, the proportion of methylated sites was relatively continuous. The mean percentage of methylated RRACH sites was 61.1% (median = 65.0%), indicating that approximately two-thirds of all RRACH sites are methylated on average.

To explore whether RRACH density influences methylation proportion, we analyzed subsets of genes with high and low RRACH counts, as defined in the DAVID GO analysis. Among the 1,000 genes with the highest number of RRACH sites, the mean methylation level was 64.7% (N = 832). In contrast, among the 1,000 genes with the lowest number of RRACH sites, a higher mean methylation level of 70.9% was observed (N = 699).

A small number of genes displayed apparent methylation percentages exceeding 100%, which is biologically implausible. This likely results from two factors. First, m^6^A methylation can occur at DRACH motifs, where D represents A, G, or U (Huang et al., 2020). Because our algorithm only counts canonical RRACH sites, it underestimates the total number of potential methylation sites, which can produce inflated percentages. Second, some RMBase datasets include cancer-derived cell lines; as explored later in this study, cancer-associated mutations can both increase and decrease the number of RRACH motifs, potentially leading to overestimation of methylation frequency.

### 3.5 Selective pressures on RRACH motifs associated with m6A methylation

To investigate the evolutionary relevance of RRACH motif abundance, we compared mammalian genes enriched or depleted in RRACH motifs with their orthologs in Caenorhabditis elegans, a species lacking the m^6^A methylation pathway (notably, it does retain rRNA methylation, Sendinc et al., 2020). We analyzed 500 human genes that were enriched in RRACH motifs and 500 genes that were depleted relative to the genomic background.

We observed a striking reciprocal pattern: RRACH-enriched mammalian genes had C. elegans orthologs with markedly lower RRACH motif frequencies (Wilcox test, p< 2.2e-16), whereas RRACH-depleted mammalian genes corresponded to C. elegans orthologs with higher RRACH frequencies (Wilcox test, p< 2.2e-16; Figure 3). This pattern suggests that, in the presence of an active m^6^A pathway, selection may act to maintain or reduce methylation motifs in a gene-specific manner. In contrast, in organisms lacking m^6^A methylation, such motifs may evolve neutrally or under distinct selective constraints. Overall, these findings support the view that RRACH motif abundance is shaped by selective pressures associated with the functional roles of the m^6^A methylation machinery.

**Figure 3.**
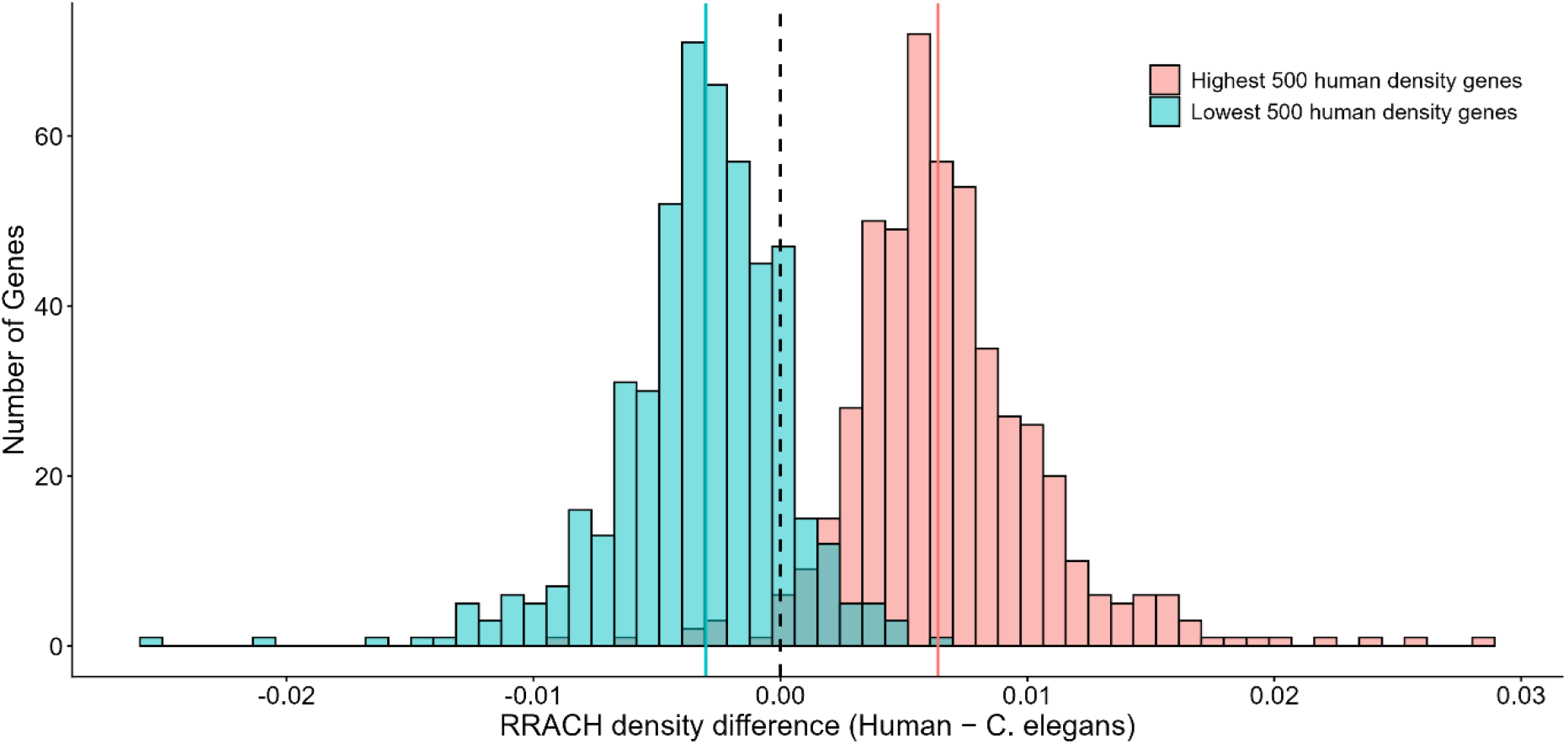
Reciprocal RRACH motif abundance in *C. elegans* orthologs is consistent with selective pressures associated with m^6^A methylation. Distributions show the difference in RRACH motif density (human minus *C. elegans*) for genes that are RRACH-enriched (red, top 500) or RRACH-depleted (blue, bottom 500) in humans. Positive values indicate higher RRACH density in humans, whereas negative values indicate higher density in *C. elegans*. Solid colored lines indicate group medians, and the dashed vertical line marks zero difference.

### 3.6 Codon Usage Bias in the Formation of RRACH Sites

Unlike CpG codon dyads analyzed previously (Creasey and Tauber, 2024), the number of possible codons that can give rise to RRACH motifs is considerably larger. At least 22 amino acid pairs can generate an RRACH site via specific codon combinations, excluding the greater number of possibilities involving codon triads.

To determine whether specific amino acid dyads are more likely to appear in RRACH-forming codons than expected, we calculated the frequency of each dyad across coding sequences and compared the number of RRACH-compatible codons to those that do not form the motif. Frequencies were normalized based on the number of codon combinations capable of generating RRACH motifs for each pair.

As shown in Figure 4, several amino acid pairs display strong biases toward either enrichment or depletion of RRACH-forming codons. In the absence of codon usage bias, the frequencies of RRACH and non-RRACH codons would be similar, resulting in bars of equal height. This is observed for some pairs, such as Arg–Leu and Arg–Pro, but most pairs deviate from this expectation.

**Figure 4.**
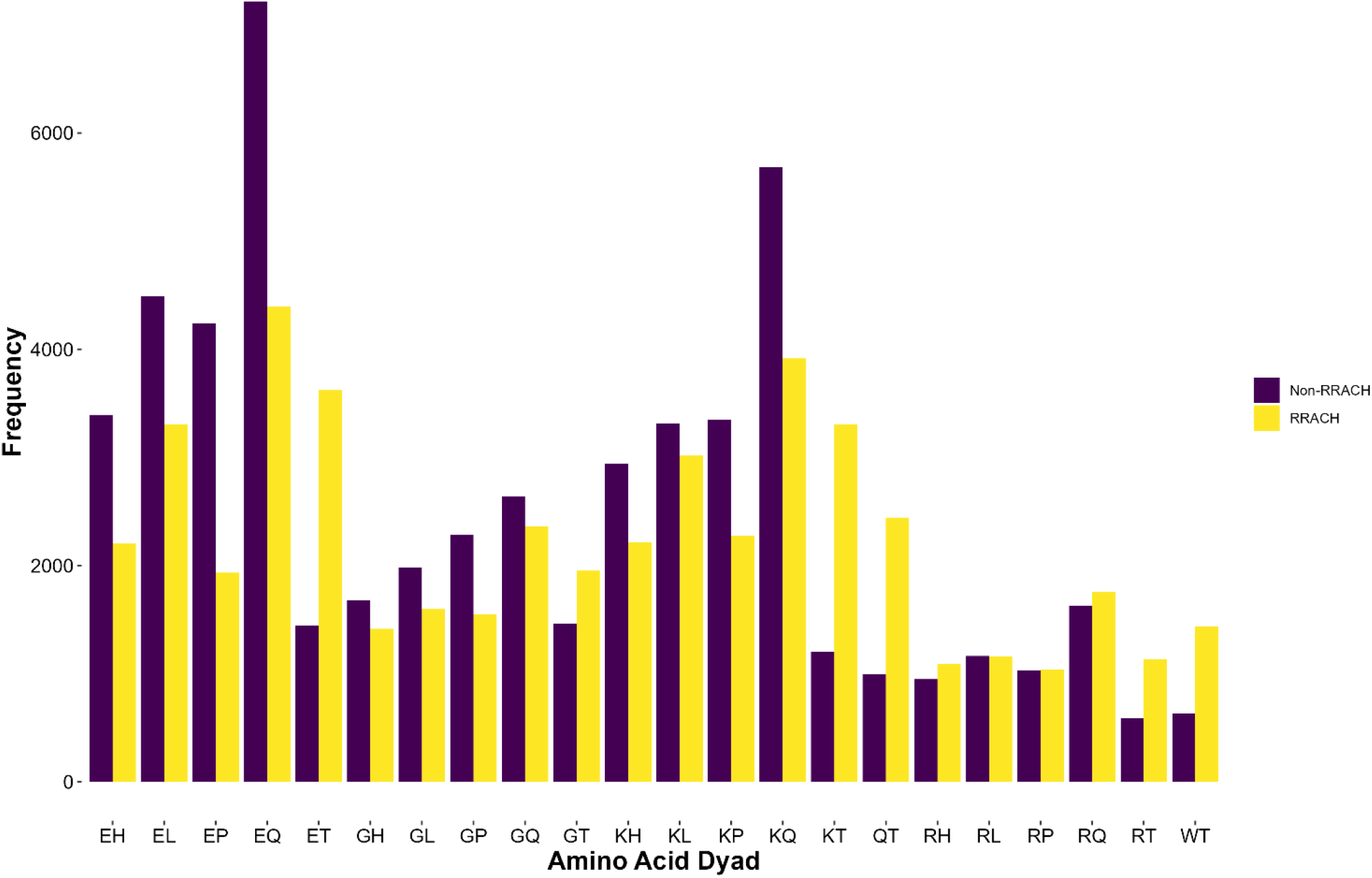
Normalized frequency of RRACH-forming (yellow) and non-RRACH (purple) codon variants for each amino acid pair.

To quantify the statistical significance of these deviations, we performed a χ^2^ test for each amino acid pair (Table 3). Many pairs showed significant depletion of RRACH motifs, including Glu–Pro, Gly– Pro, and Lys–Leu. Conversely, several pairs, particularly those ending in threonine, showed strong and highly significant enrichment. All amino acid pairs ending in threonine were significantly enriched for RRACH-forming codons, with additional enrichment observed for Arg–Gln and Arg– His.

**Table 3.** Representative amino acid pairs showing enrichment or depletion of RRACH-forming codons. Full results for all pairs are provided in Supplementary Table S1.

| Amino acid pair | Change | $\chi^2$ | *p*-value |
| --- | --- | --- | --- |
| Glu–Pro | Depletion | 1786.51 | $<1 \times 10^{-300}$ |
| Gly–Pro | Depletion | 423.51 | $4.2 \times 10^{-94}$ |
| Lys–Leu | Depletion | 37.05 | $1.2 \times 10^{-9}$ |
| Arg–Pro | Neutral | 0.16 | 0.69 |
| Arg–Gln | Enrichment | 7.55 | $6.0 \times 10^{-3}$ |
| Arg–His | Enrichment | 16.41 | $5.1 \times 10^{-5}$ |
| Arg–Thr | Enrichment | 1064.95 | $1.4 \times 10^{-233}$ |
| Lys–Thr | Enrichment | 1520.83 | $<1 \times 10^{-300}$ |

All amino acid pairs showing the most significant enrichment for RRACH sites end with threonine, although Arg–Gln and Arg–His also show positive selection. Most other pairs exhibit depletion of RRACH sites, suggesting that threonine-based RRACH motifs may represent a preferred structural or functional context for m^6^A deposition.

Beyond dyads, RRACH motifs can also be generated by codon triads (NNR–RAC–HNN). Triads are inherently more flexible because of variation in the third redundant position and the presence of the variable H base at the conserved site. Consequently, many more codon combinations (421 compared to 22 for dyads) can potentially generate an RRACH motif while maintaining the encoded amino acid sequence.

Using the same analytical approach, we extended our analysis to triads. Of the 421 different triads tested, only one showed no significant difference between non-RRACH and RRACH triads. Of the remaining 420, 325 showed significant overrepresentation of RRACH sites, and 95 showed significant depletion. Overall, this suggests that triads are more frequently used to generate RRACH sites where they are functionally required.

### 3.7 Gain and loss of RRACH sites associated with cancer

Mutation datasets, pre-split into cancer-related genes and all genes with detected mutations, were obtained from the COSMIC database. The two datasets yielded broadly similar results, even for genes appearing in both sets. To assess consistency between datasets, we conducted a correlation test and found a strong concordance in loss and gain rates of RRACH sites (Figure 5; *r* = 0.967, *P* < 0.001).

**Figure 5.**
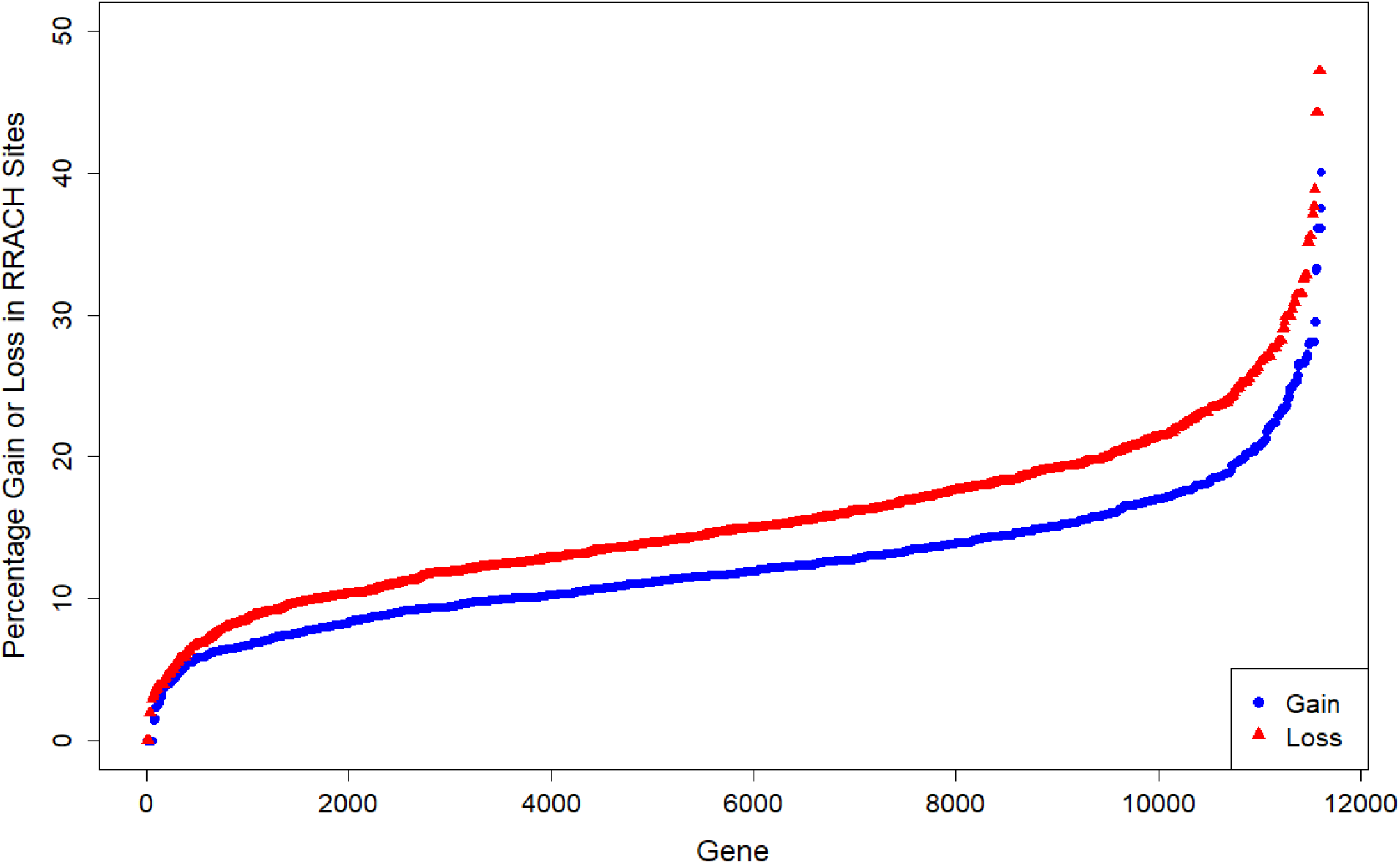
Percentage of mutations that lead to loss (red) or gain (blue) of RRACH sites as a proportion of total mutations per gene in the core cancer gene set.

From these data, we determined that 4.6% of mutations in the core cancer gene dataset led to the loss of an RRACH site, while 3.6% led to the gain of one (in the total dataset, the respective values were 4.4% and 3.6%). When analyzed gene by gene, a wide range of RRACH loss or gain percentages was observed (Figure 5). Each point in Figure 5 represents a gene, showing the percentage of mutations that lead to either the loss (red) or gain (blue) of RRACH sites. Both distributions approximate normality, with loss events generally more frequent than gains.

To examine tissue-specific patterns, we separated the dataset by organ of origin and quantified, for each cancer type, the per-gene rate at which mutations produced the loss or gain of an RRACH site (normalised per gene as defined in Methods; Figure 6). The comparison revealed considerable tissue-specific variation in RRACH site dynamics. Among the cancer types analysed, skin showed the strongest bias toward RRACH loss and cervix the strongest bias toward gain, whereas large intestine showed the highest absolute gain rate among the significant tissues. By contrast, several cancer types, including breast and pancreatic cancer, showed no significant imbalance between loss and gain (two-sided paired Wilcoxon signed-rank tests with Benjamini–Hochberg correction; Figure 6; per-tissue rates, sample sizes, and adjusted *p*-values in Supplementary Table S2), indicating that the balance between RRACH loss and gain differs across tissues and tumour types. Notably, although skin showed the strongest bias toward RRACH loss, it did not have the highest absolute loss rate; rather, its distinctiveness arises from a relative scarcity of gain events, directly visible in Figure 6 as an unusually low gain value.

**Figure 6.**
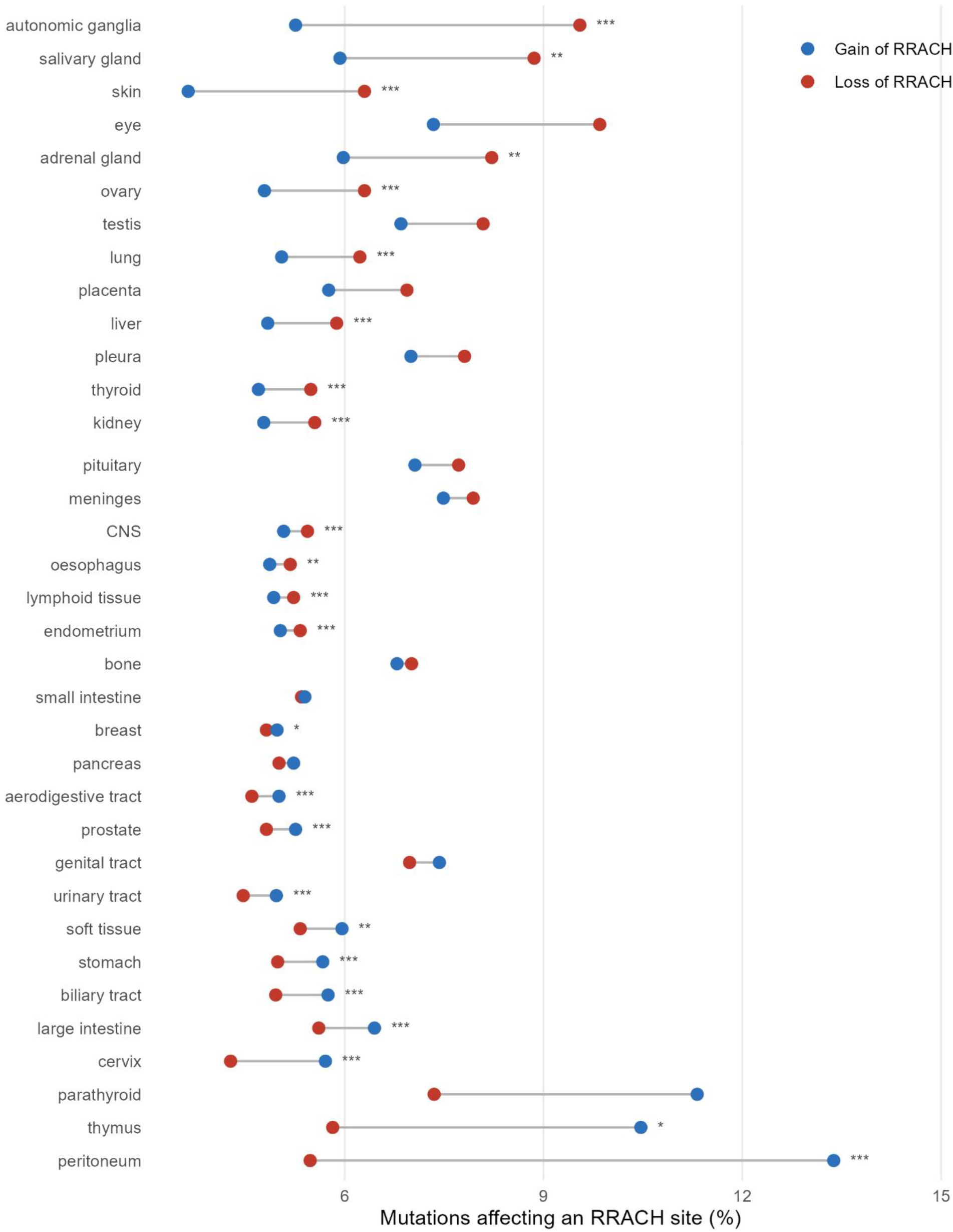
Tissue-specific loss and gain of RRACH sites in cancer. Somatic mutations in the COSMIC core cancer gene set were obtained from COSMIC (Sondka et al., 2024). For each gene, the rate of RRACH loss (red) and gain (blue) was computed as the number of somatic mutations destroying or creating an RRACH site, normalised by the geometric mean of the gene’s total mutation count and its number of RRACH sites; points show the per-gene mean for each cancer type (×100), and the grey bar spans the loss–gain difference. Cancer types are ordered by net bias (loss − gain), from most loss-biased (top) to most gain-biased (bottom). Within each cancer type, per-gene loss and gain rates were compared using a two-sided paired Wilcoxon signed-rank test, with Benjamini– Hochberg correction across the 27 cancer types shown; asterisks denote the adjusted significance (*** P < 0.001, ** P < 0.01, * P < 0.05), and unmarked cancer types are not significant. Cancer types represented by fewer than 150 transcript records were excluded. Skin shows the strongest bias toward RRACH loss and cervix the strongest bias toward gain, while large intestine has the highest absolute gain rate, while many tissues show no significant imbalance, consistent with context-dependent remodelling of methylation potential. Exact loss and gain rates, the number of gene records per cancer type, and the raw and Benjamini–Hochberg-adjusted p-values underlying the significance markers are provided in Supplementary Table S2.

These tissue-specific differences were corroborated by a χ^2^ contingency test comparing the rates of RRACH site loss and gain across cancer types (Figure S1). Of the 29 organs analyzed, 17 showed a significant difference between loss and gain. Because χ^2^ contingency tests can be disproportionately influenced by categories with large residuals, we performed a sensitivity analysis excluding cancer types with extreme contributions to the test statistic, such as skin and large intestine. When these categories were removed, several additional cancer types, including esophageal cancer, showed significant differences in RRACH site loss versus gain (standardized residual increased from –0.14 to 4.35, *p* < 0.001), indicating that strong deviations in a few cancer types can obscure subtler patterns in other tissues. As these are paired comparisons, a high significance value reflects a large relative difference between loss and gain rather than an absolute increase in either category.

There was no consistent pattern across cancer types: some showed significant loss or conservation of RRACH sites, whereas others exhibited gains or selection against gain. This heterogeneity aligns with previous reports that the role of m^6^A methylation in tumorigenesis is highly context dependent, reflecting differences among tissues and cancer lineages in methylation dynamics and regulatory mechanisms.

## 4 Discussion

This study provides a large-scale comparative analysis of RRACH motif distribution across mammalian coding sequences and its relationship to m^6^A RNA methylation. By integrating cross-species conservation data, methylation site annotation, and codon usage analyses, we demonstrate that RRACH abundance is non-random, evolutionarily constrained, and functionally associated with specific gene classes. These results suggest that selection acts on synonymous sites to balance the potential for RNA methylation against competing pressures such as DNA methylation and codon usage bias.

Using *Caenorhabditis elegans*, a species lacking canonical mRNA based m^6^A methylation (Sendinc et al., 2020), as a comparative reference, we observed reciprocal differences in RRACH motif density between mammals and nematodes. Mammalian genes enriched in RRACH density often have C. elegans orthologs with reduced motif density, while mammalian genes depleted of RRACH density correspond to orthologs with higher frequencies. This pattern indicates that, in the presence of an active m^6^A pathway, selective forces may act to maintain or eliminate RRACH sites in a gene-specific manner. In species lacking this pathway, RRACH motifs likely evolve neutrally or under unrelated sequence constraints.

One of the more striking findings of this work is the reciprocal positional pattern of RRACH and CpG motifs within coding regions. RRACH sites were depleted near the 5′ end of genes and became evenly distributed beyond roughly one-third of the coding sequence, whereas CpG codon dyads, associated with DNA methylation, tend to occur closer to the 5′ end (Creasey & Tauber, 2024).

Because RRACH motifs cannot contain CpG dinucleotides, these two methylation-related signals are mutually exclusive. This relationship suggests that DNA and RNA methylation systems impose opposing constraints on sequence composition, effectively partitioning the coding region into zones of preferred methylation potential. Statistical correlations between RRACH and CpG motif distributions support this interpretation, revealing a negative relationship in the 5′ portion of genes and a positive one toward the 3′ end. Such complementary distributions imply that selection acting at both the DNA and RNA levels has co-shaped coding sequence evolution.

Functional enrichment analysis showed that genes with high RRACH density were particularly enriched for ubiquitin-like conjugation and cell cycle–related processes. Both are characterized by tightly regulated post-transcriptional control, and the high occurrence of RRACH motifs in these genes may facilitate dynamic transcript regulation through m^6^A modification (Zuin et al., 2014). By contrast, genes depleted of RRACH motifs were enriched for transmembrane proteins and HOX or homeobox genes. The absence of RRACH sites in HOX genes is likely a byproduct of their CpG-rich coding regions (Creasey & Tauber, 2024), given that CpG sequences cannot coexist within RRACH motifs. The association with transmembrane proteins is less easily explained but may reflect their constitutive expression and limited need for post-transcriptional regulation.

The distribution of methylated RRACH sites across genes was consistent with previous findings that m^6^A is more abundant near the 3′ end of coding sequences (Meyer et al., 2012). Our analysis refines this pattern, showing that depletion is confined to the first 20–30% of gene length, beyond which methylation potential remains stable. A recent study by Uzonyi et al. (2023) proposed an “exclusion model” of m^6^A deposition, in which methylation is inhibited near exon–intron junctions but otherwise widespread. Because terminal exons are typically longer, the apparent 3′ enrichment may simply reflect the greater availability of methylation-permissive regions. The relatively uniform presence of RRACH motifs along the remainder of the coding sequence supports this idea and suggests that RRACH sites maintain potential for methylation throughout the transcript body.

We estimated that roughly two-thirds of RRACH motifs are methylated, though this likely underestimates the true rate. m^6^A is dynamic, tissue-specific, and temporally regulated, meaning that transient or context-dependent modifications are often missed. Indeed, comparison with earlier versions of the RMBase dataset shows that the number of identified methylated sites has roughly doubled between versions 2.0 and 3.0 (Xuan et al., 2018). This likely reflects improvements in detection methods rather than true evolutionary change. Some overestimation is possible due to inclusion of cancer-derived data, where mutations can both create and remove RRACH motifs.

However, these biases do not affect the overall conclusion that RRACH motifs are widespread and frequently methylated in mammalian genes.

A particularly notable result was the discovery of codon usage bias affecting RRACH formation. Of the 22 amino acid dyads capable of generating RRACH motifs, only two showed no significant bias, while the remainder exhibited either enrichment or depletion. Dyads ending in threonine were strongly enriched for RRACH motifs. Although this is partly explained by threonine’s codon structure (ACN), enrichment persisted even after accounting for codon combination frequency. The likely explanation is avoidance of the CpG-containing threonine codon ACG, which is selectively suppressed in mammals (Li & Zhang, 2014). This avoidance indirectly favors codons that contribute to RRACH formation. A similar pattern was observed for arginine–histidine and arginine–glutamine pairs: arginine codons beginning with CG are underrepresented due to CpG suppression, while AG-starting codons, which can form RRACH motifs, are more common. These examples highlight how DNA-level methylation constraints and codon usage biases interact to shape RNA methylation potential. When adjusted for CpG suppression, the overall trend indicates a mild selection against RRACH motifs, suggesting that they are maintained only where functionally beneficial.

A complementary line of evidence reinforces the inference that m^6^A-associated sequence features are under selection: across 447 mammalian species, m^6^A sites with high methylation stoichiometry are more evolutionarily conserved than weakly methylated sites, consistent with purifying selection preserving functionally important sites (Pike & Schwartz, 2026). Our results address a different aspect of the same problem. Rather than asking whether existing methylated sites are conserved, we ask how synonymous codon choice biases the formation and avoidance of RRACH motifs in the first place, and we find patterns consistent with selection acting in both directions, favouring motif formation in some gene classes and disfavouring it in others. Together, these observations suggest that selection shapes m^6^A potential at two levels: the retention of high-value sites once established, and the codon-level tuning of where such sites can arise

The analysis of cancer mutation datasets revealed further complexity in the regulation of RRACH motifs. Different cancers displayed distinct patterns of motif gain and loss, with no universal direction of change. Skin cancers showed a relative excess of RRACH site loss, whereas large intestine cancers showed gains. Lung cancers exhibited frequent RRACH loss despite upregulation of methyltransferase “writer” enzymes (Wanna-udom et al., 2020), suggesting compensatory sequence evolution to balance excessive methylation. Such remodelling may have functional consequences: the loss of RRACH sites in tumour suppressor genes could reduce m^6^A-mediated transcript decay, while gains in oncogenes could destabilize transcripts. These findings support a model in which cancer-associated mutations dynamically reshape local methylation potential, contributing to altered RNA metabolism in tumours.

The remodelling of RRACH content by cancer-associated mutations that we observe builds on growing evidence that somatic mutations frequently alter m^6^A sites in human tumours. Chen et al. (2024) documented the prevalence of synonymous mutations affecting m^6^A modification sites across human cancers, and Lan et al. (2025) went further to show that such synonymous mutations can promote tumorigenesis specifically by disrupting m^6^A-dependent mRNA metabolism, identifying discrete m^6^A-disrupting mutations enriched in tumour suppressors such as CDKN2A and BRCA2 and validating their effects on transcript stability by epitranscriptomic editing. Our analysis complements these site- and gene-focused studies by describing the broader, tissue-resolved landscape of RRACH gain and loss across cancer types. Whereas previous work has largely emphasised the disruption (loss) of m^6^A sites, our data resolve both the loss and the gain of RRACH motifs and reveal tissue-specific asymmetries between them, placing these mutational effects in a genome-wide and comparative context. The convergence of these approaches (prevalence surveys, single-site mechanistic validation, and our cross-tissue statistical analysis) supports the view that the somatic remodelling of RRACH motifs is functionally consequential for m^6^A regulation rather than reflecting neutral mutational turnoverCollectively, our results reveal that RRACH motif abundance and distribution are shaped by overlapping layers of selection acting at both the nucleotide and functional levels. Codon composition, CpG suppression, gene function, and cellular context each contribute to the observed variation. This interplay between genetic and epigenetic constraints suggests that synonymous sites are far from neutral, instead encoding information relevant to both DNA and RNA methylation systems. We propose that these systems jointly define an “epigenetic coding layer” within protein-coding sequences, integrating transcriptional and post-transcriptional regulation.

Future work should extend these analyses beyond mammals to other vertebrates and invertebrates with independently gained or lost m^6^A pathways, allowing direct tests of evolutionary hypotheses about methylation-related sequence selection. At a mechanistic level, recent advances in nanopore sequencing now enable direct detection of m^6^A at single-molecule resolution (Teng et al., 2024), offering opportunities to quantify site-specific methylation dynamics across developmental stages and tissues. Such studies will help clarify how sequence-level constraints translate into regulatory flexibility and how the evolution of RRACH motifs contributes to gene expression control in both health and disease.

## 6 Author Contributions

LDC: Investigation, writing (original draft), ET: Conceptualisation, Supervision and Writing (review and editing).

## 20 Data Availability Statement

The datasets generated and analyzed during this study have been deposited in Zenodo and will be made publicly available upon acceptance of the manuscript.

